# Brain Structure Shapes Function through higher-order Functional Interactions

**DOI:** 10.64898/2026.06.22.733911

**Authors:** Shichao Su, Mingrui Zhuang, Marco Palombo, Manhua Liu, Xi Jiang, Tuo Zhang, Hongkai Wang, Songyao Zhang

**Affiliations:** Central Hospital of Dalian University of Technology, Dalian Municipal Central Hospital, Dalian, China; School of Biomedical Engineering, Faculty of Medicine, Dalian University of Technology, Dalian, China; Brain Research Imaging Centre, Cardiff University, Cardiff, Wales, United Kingdom; Institute of Artificial Intelligence, Shanghai Jiao Tong University, Shanghai, China; The Clinical Hospital of Chengdu Brain Science Institute, MOE-K Lab for NeuroInformation, Brain-Apparatus Communication Institute, University of Electronic Science and Technology of China, Chengdu, China; School of Automation, Northwestern Polytechnical University, Xi’an, China; Liaoning Key Laboratory of Integrated Circuit and Biomedical Electronic System

**Keywords:** Higher-order functional interactions, Structural-functional constraint strength, Multimodal neuroimaging, Cognitive phenotype, ***O***-information

## Abstract

Brain function is deeply embedded within multiscale structural architecture. Conventional studies predominantly utilize pairwise connectivity networks to investigate structure-function relationships. However, this low-dimensional perspective overlooks multi-region collaborations for complex cognition. Consequently, whether and how anatomy constrains such higher-order functional networks remains unresolved. To address this pivotal question, we utilize an information-theoretic ***O***-information approach to characterize higher-order functional interactions (HOIs). By reconstructing individual-level HOIs from multimodal structural networks, we directly validate the structural constraint on HOIs. The resulting reconstruction coefficients are defined as structural-functional constraint strength (SFCS), serving as a quantitative vehicle to decipher how anatomy shapes these higher-order networks. SFCS uncovers a highly heterogeneous structural constraint landscape across data modalities, spatial regions, and informational interaction modes. Crucially, individualized SFCS robustly predicts multi-domain cognitive phenotypes, showing higher sensitivity for informant-reported than patient-reported assessments. Finally, we show that this landscape undergoes pathological, mode-specific reorganization in Alzheimer’s disease. Cross-scale alignment with spatial transcriptomics further demonstrates that this macroscale network remodeling is coupled with microscale metabolic and regulatory gene pathways. Collectively, our findings not only validate the structural constraint on higher-order functional networks but also decipher its precise underlying mechanisms. This constraint paradigm plays a pivotal role in shaping diverse cognitive capabilities, while its pathological disruption in Alzheimer’s disease highlights the potential of SFCS as a biomarker for tracking neurodegenerative network impairments.

## 1 Introduction

Brain function does not emerge independently of anatomy but is embedded within the brain’s multiscale structural architecture [1, 2]. Accordingly, understanding the relationship between brain structure and function has long been a central question in systems neuroscience [3, 4]. Extensive evidence suggests that brain function and cognitive performance are governed by a multiscale structural hierarchy, ranging from macro-scale white matter connectivity and cortical morphology to micro-scale neuronal organization [5, 6]. Network theory provides a parsimonious and powerful framework for investigating how brain function is embedded within the brain’s structural organization [7]. Although network-based analyses have revealed principles of structure-function coupling and enabled bidirectional prediction between structural and functional brain networks, these studies have predominantly focused on pairwise interactions, largely overlooking complex interactions among multiple regions [2, 8]. As a result, current understanding of structure-function relationships remains largely confined to low-order functional interactions.

Brain information processing is inherently distributed and cooperative, with coordinated activity among multiple regions jointly shaping the brain’s functional state during diverse cognitive tasks and spontaneous activity [9]. Consequently, pairwise connectivity alone may be insufficient to fully characterize the complexity of functional brain organization [10]. To bridge this gap, an increasing number of studies have recently employed higher-order brain network models to characterize brain function, including hypergraphs that explicitly encode multi-region relationships [11, 12], simplicial complexes that capture topological dependencies beyond pairwise connections [13], and deep learning frameworks that learn latent higher-order interaction patterns from neuroimaging data [14]. Such studies have demonstrated that higher-order brain networks can reveal important organizational features that are difficult to capture using traditional pairwise networks, including collective dependencies among multiple regions and distributed information integration [15]. Therefore, investigating brain functional organization from the perspective of higher-order interactions is increasingly recognized as an important direction for understanding complex brain dynamics.

Although methods such as hypergraphs and deep learning can successfully characterize higher-order architectures, they often rely on specific model assumptions and struggle to directly quantify the information processing mechanisms underlying these complex connections [16, 17]. Information-theoretic approaches, particularly those based on ***O***-information, provide a principled and model-independent framework for quantifying higher-order interactions among multiple brain regions [9]. Unlike traditional methods, ***O***-information can directly capture the joint dependency structure of a multivariate system and decompose it into two fundamental forms: redundancy and synergy [18]. Redundancy reflects information that is shared across multiple brain regions, whereas synergy refers to information that emerges only from the joint state of multiple regions. This decomposition provides an important perspective for understanding how distributed neural systems balance information sharing and information integration [16]. Consequently, this information-theoretic decomposition offers a systematic and interpretable framework for investigating higher-order functional organization in the brain.

While ***O***-information allows us to capture the dynamics of higher-order functional networks, its underlying anatomical basis remains completely unexplored. Although a general correspondence between brain structure and function is well-recognized, existing evidence mostly stops at statistical correlations [19]. Unlike simple pairwise correlations, redundancy and synergy define the fundamental informational logic governing how signals are integrated across multiple regions [9]. Therefore, a critical unanswered question is whether the brain’s physical architecture—specifically, structural connectivity and morphometric similarity—actively constrains the spatial heterogeneity of these higher-order informational states [20]. Unraveling this mechanistic relationship may not only illuminate the structural basis of brain functional organization and cognitive variability but also provide novel insights into the mechanisms underlying neurological disorders.

In this study, we propose an information-theoretic framework for higher-order structure–function analysis to investigate whether and how brain structure constrains higher-order functional interactions. Specifically, we constructed structural connectivity (SC) from diffusion MRI and morphometric similarity networks (MSNs) from T1-weighted imaging, and derived higher-order interactions (HOIs) from functional MRI, which were further decomposed into redundant and synergistic components. Based on these representations, we developed a structure-driven reconstruction model to test whether SC and MSNs can reconstruct the spatial organization of higher-order functional interactions at the individual level. We further interpreted the model parameters as indicators of the strength with which brain structure constrains higher-order information organization, allowing us to systematically examine the roles of different brain regions and structural modalities in constraining redundant and synergistic interactions. Finally, we investigated how these structural constraint patterns relate to cognitive phenotypes and functional alterations associated with Alzheimer’s disease (AD), and combined enrichment analysis and meta-analytic decoding to explore their potential neurobiological underpinnings.

## 2 Results

To characterize how structural architecture constrains higher-order functional interactions, we established a multimodal modeling framework (Fig. 1) across 301 individuals (510 scans, Table S1). Higher-order functional interactions (HOIs) were quantified using third-order ***O***-information computed from fMRI signals (see Methods 5.1). The sign of the ***O***-information distinguishes two interaction regimes with distinct network implications: positive values indicate redundancy-dominant interactions, whereas negative values indicate synergy-dominant interactions. To focus on the most representative and directionally unambiguous informational states, we extracted the upper 20% and the lower 20% of the HOI distribution for each participant to represent the core “redundancy” and “synergy” components, respectively (Fig. 1B left). This thresholding isolates the core functional architecture by mitigating the influence of weak, mixed dependencies near zero. Detailed validation for this specific cutoff choice is provided in Supporting Information (see Fig. S1, S2)

**Fig. 1:**
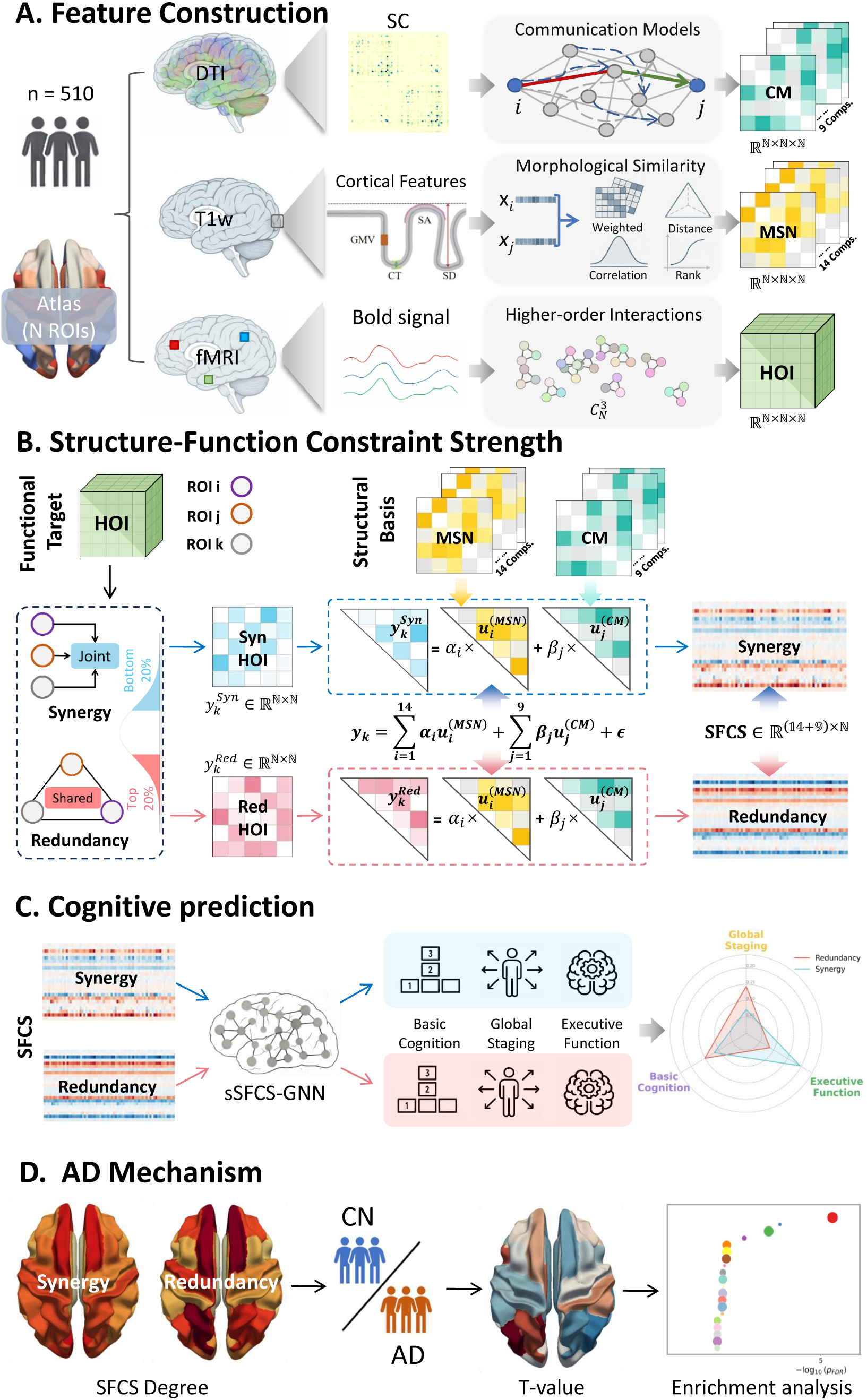
Overview of the analysis framework. (A) Feature construction from T1-weighted MRI, diffusion MRI, and resting-state fMRI. This process yields a comprehensive topological basis set comprising 14 morphometric similarity networks (MSNs) and 9 structural connectivity (SC) matrices, alongside three-dimensional tensors of higher-order functional interactions (HOIs). (B) Structure-driven GLM reconstruction. The synergy (bottom 20%) and redundancy (top 20%) HOI slices (**y***_k_*) are fitted using the 23 structural components (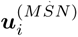 and 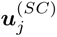) to extract distinct structure-function constraint strength (SFCS) profiles. (C) Cognitive prediction across distinct domains using a graph neural network (sSFCS-GNN) driven by subject-specific SFCS profiles. (D) Clinical and biological characterization, encompassing AD-related alterations in regional SFCS degree and associated gene enrichment analysis.

Crucially, to match the inherent complexity of the triadic functional landscape, we moved beyond the limitations of isolated structural connectivity (SC) or morphometric similarity networks (MSNs). Instead, we constructed a comprehensive structural basis set comprising 23 distinct structural components (Table 1 and Methods 5.2), as illustrated in Fig. 1A. This ensemble integrates diverse anatomical principles-ranging from physical routing and communication efficiency to morphological dependencies, thereby providing a more comprehensive brain-structural informational representation for the characterization of higher-order functional interactions. Group-level analysis revealed that these 23 structural components are largely orthogonal to one another, exhibiting minimal cross-modal redundancy and strong informational complementarity, which establishes an optimal basis set for the subsequent HOI reconstruction (Fig. S3). As shown in Fig. 1B, within a general linear modeling (GLM) framework, we fitted each slice (corresponding to specific brain regions) of the three-dimensional HOI tensor using these 23 structural representations, yielding separate Structure-Function Constraint Strength (SFCS) profiles for redundancy and synergy (see Methods 5.3). This unified representation then served as the foundation for subsequent analyses of cognitive prediction (Fig. 1C), AD mechanism and gene-expression associations (Fig. 1D).

**Table 1:** Summary of the 23 structural components used in the study.

| Category | Subcategory | Full name | Abbrev. (short) |
| --- | --- | --- | --- |
| Morphological similarity | Rank-based Correlation | Spearman’s rank correlation | Spearman |
|  |  | Kendall’s rank correlation | Kendall |
|  |  | Squared Spearman correlation strength | Spearman-sq |
|  |  | Squared Kendall correlation strength | Kendall-sq |
|  | Distance-based Dissimilarity | Euclidean distance | Euclidean |
|  |  | Cityblock (Manhattan) distance | Cityblock |
|  |  | Chebyshev distance | Chebyshev |
|  |  | Cosine distance | Cosine |
|  | Weighted Dissimilarity | Canberra distance | Canberra |
|  |  | Bray–Curtis dissimilarity | Bray-Curtis |
|  | Generalized Correlation | Distance correlation (biased estimator) | dCor-biased |
|  |  | Distance correlation (unbiased estimator) | dCor |
|  |  | Hilbert–Schmidt independence criterion (biased estimator) | HSIC-biased |
|  |  | Hilbert–Schmidt independence criterion (unbiased estimator) | HSIC |
| Communication model | Topological Distance | Weighted shortest path length | SPL |
|  |  | Number of hops along shortest path | Hops |
|  |  | Predecessor matrix of shortest path | Pmat |
|  |  | Binary distance | BD |
|  | Topological Similarity | Matching index | MI |
|  |  | Path transitivity | PT |
|  | Communicability and Dynamical | Communicability | F |
|  |  | Mean first passage time | MFPT |
|  |  | Flow graph dynamics | FGD |

### 2.1 Multimodal structural components accurately reconstruct higher-order functional interactions

We first tested whether multimodal structural representations could accurately reconstruct HOIs. Using a general linear model built on 23 two-dimensional structural matrices, reconstructed networks showed consistently high agreement with real HOIs across 510 scans from 301 individuals and all region-defined slices. Quantified by the Pearson correlation between real and reconstructed HOIs, most slices in both hemispheres were stably distributed within the 0.8-0.85 range, with broadly similar distributions across regions (Fig. 2A). These results indicate that, despite the high dynamicity and complexity of brain functions, multimodal structural representations reliably capture the higher-order functional interactions at the slice level. This robust reconstruction performance was visually corroborated in representative regional slices. Example heatmaps, displaying reconstructed values in the upper triangles and real values in the lower triangles, demonstrated a striking visual correspondence (Fig. 2B). Moving from slice-level summaries to the complete triadic architecture, we examined the element-wise accuracy of the full 3D HOI tensor in an example subject. The reconstructed and real patterns exhibited substantial correlations for both synergy (*r* = 0.476*, n* = 10480) and redundancy (*r* = 0.533*, n* = 10481) components (Fig. 2C). The high density core regions (warm-colored) tightly clustered around their respective regression lines. Therefore, despite the high dimensionality and vast number of statistical elements inherent in full HOIs, structure-constrained models still achieve robust reconstruction performance and maintain remarkably strong correlation scores for both higher-order informational processes.

**Fig. 2:**
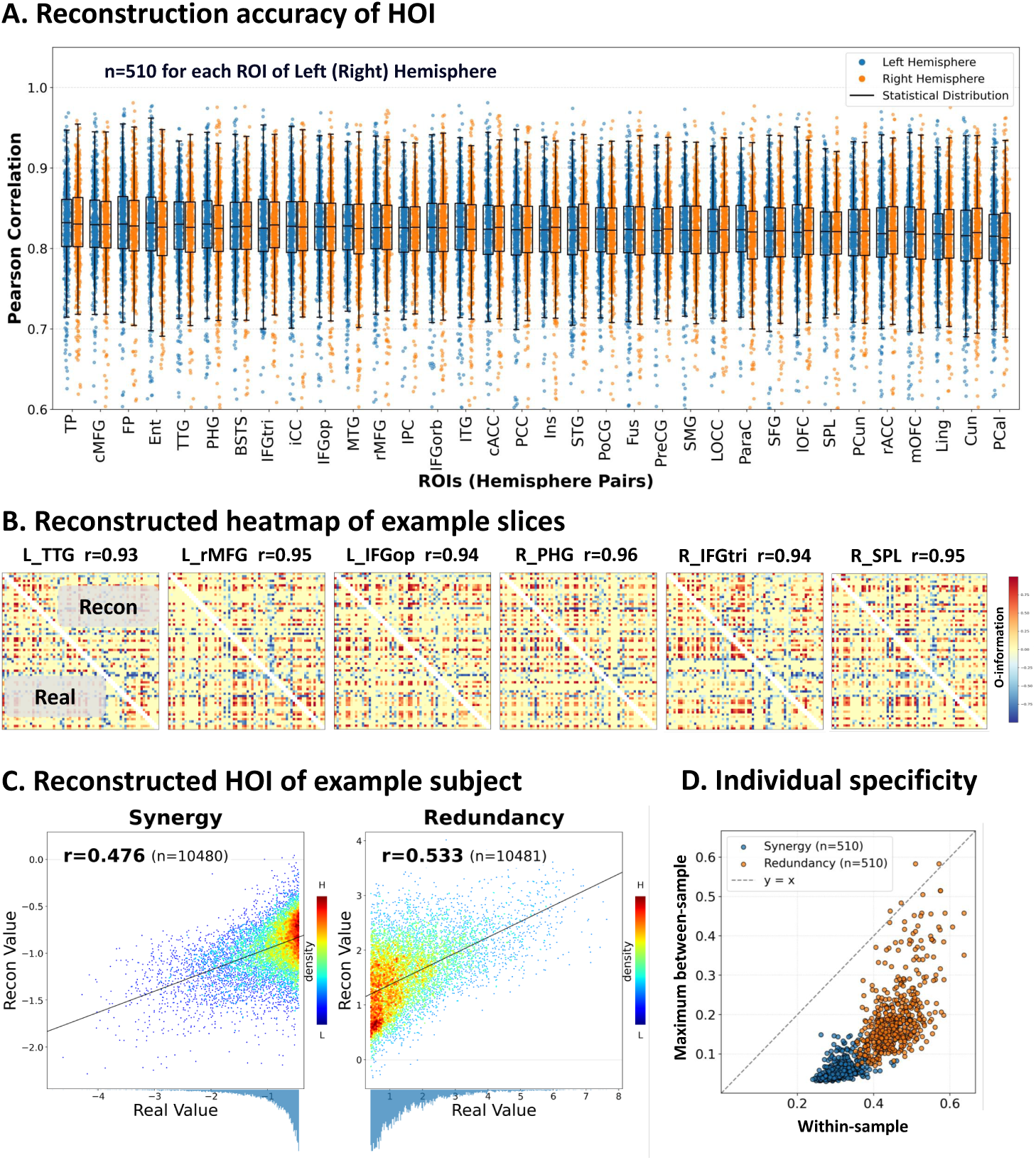
Slice-wise reconstruction of higher-order functional interactions (HOIs). (A) Pearson correlations between real and reconstructed HOIs across all region-defined slices from 510 scans acquired in 301 individuals. Each point represents the reconstruction accuracy of one region-defined slice from a single scan, which was treated as an independent sample in the analysis; blue and orange denote slices from the left and right hemispheres, respectively. Brain regions are ordered from high to low according to the mean of the median reconstruction accuracies across the two hemispheres. (B) Representative heat maps of reconstructed and real HOIs for example slices. (C) Scatter plots comparing real and reconstructed synergistic and redundant HOIs in an example sample; warm-to-cool colors indicate high-to-low local point density. (D) Individual-specificity analysis, showing within-sample similarity versus maximum between-sample similarity. Each point represents the reconstruction accuracy of the whole-brain three-dimensional redundant (orange) or synergistic (blue) HOI from a single sample.

Given the exceptional reconstruction capacity at the group and intra-subject levels, we next asked whether the reconstructed networks retained individual specificity. For both synergy and redundancy, we compared the within-sample similarity (between an individual’s reconstructed and real networks) against the maximum between-sample similarity (between that individual’s reconstructed network and the reconstructed networks of all other participants). As depicted in Fig. 2D, the scatter points for both the synergy (blue) and redundancy (orange) components fell below the identity line (*y* = *x*), indicating that within-sample similarity significantly exceeded the maximum between-sample similarity. Notably, while both modes exhibited distinct functional fingerprinting, the redundancy component generally displayed higher within-sample similarity scores than the synergy component. This finding demonstrates that the structure-constrained reconstruction not only approximates the real interaction at the group level, but also preserves individually discriminable higher-order functional features.

### 2.2 Structural components differentially constrain redundancy and synergy

The high fidelity of our HOIs reconstruction indicates that the resulting reconstruction weights capture fundamental constrain principles, offering a quantitative proxy to explore the constraint imposed by distinct structural components. Motivated by this, we sought to interpret these weights as the Structure-Function Constraint Strength (SFCS), allowing for a comprehensive group-level characterization of the structural-functional landscape across 136 cortical regions and 23 structural components. This SFCS profile revealed a stable and consistent global pattern: across all regions in both hemispheres and across most structural components, constraint strength was higher for redundancy than for synergy (Fig. 3A). This difference was evident not only as an elevation in overall magnitude, but also across feature classes, with white-matter communication models imposing generally stronger constraints than cortical morphological similarity matrices. These observations indicate that higher-order functional interactions, particularly redundancy-dominant interactions, are strongly anchored to the brain’s topological communication scaffold.

**Fig. 3:**
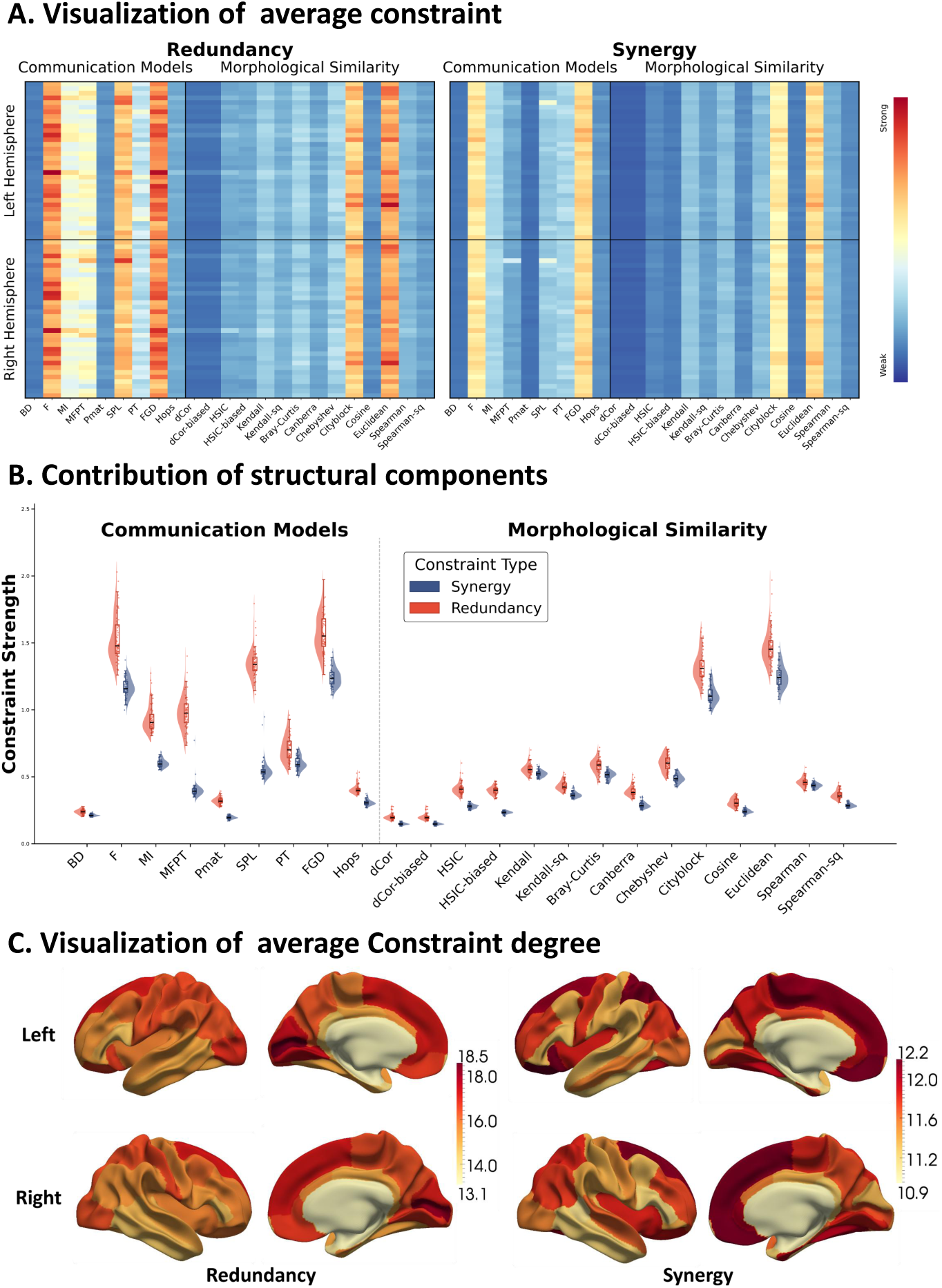
Regional and component-wise distributions of structure-function constraint strength (SFCS) for redundancy and synergy. (A) Group-average SFCS spectrum for redundancy (left) and synergy (right). Vertical axes represent 68 regions in both hemispheres, and horizontal axes represent 23 structural components categorized into communication models and morphological similarity. (B) Component-wise distributions of constraint strength across 68 cortical regions, shown separately for communication models and morphological similarity networks. (C) Cortical projections of the group-average regional SFCS degree for redundancy and synergy.

Decomposing these constraint distributions across the 68 cortical regions, this global divergence became more evident at the component level (Fig. 3B). Among communication models, redundancy and synergy showed different profiles. Most communication metrics imposed relatively stronger constraints on redundancy, whereas synergy showed a more selective dependence on a subset of communication regimes. In particular, diffusion-based communication models—notably Communicability (F) and Flow Graph dynamics (FGD)—yielded the highest constraint strengths for redundancy while remaining comparatively high for synergy. By contrast, models with moderate contributions, such as Weighted Shortest Path Length (SPL), Mean First Passage Time (MFPT), and Matching Index (MI), showed stable contributions to redundancy but smaller contributions to synergy. Low-contribution metrics (e.g., Hops, BD) showed weak effects on both. Overall, these patterns suggest that redundancy is more broadly associated with communication models, whereas synergy depends on a narrower range. The constraint profile of cortical morphological similarity was weaker overall than that of communication models, but it was not uniform. Most morphological features exhibited low-to-moderate constraint strength, whereas the Cityblock and Euclidean matrices formed the principal peaks within this class, showing comparatively stronger constraints for both redundancy and synergy. Notably, morphological similarity showed relatively parallel, albeit modest, constraints on both redundancy and synergy. These results suggest that macroscopic regional similarity may provide a broad structural background, but contributes less strongly than communication models in the present framework.

Finally, the regional distributions of constraint strength revealed distinct spatial variability patterns. As illustrated by the scatter points and violin plots in Fig. 3B, the 68 Desikan–Killiany regions exhibited a broader distribution of redundancy constraint strength than of synergy constraint strength. This pattern indicates greater inter-regional variability in the extent to which anatomical structure is associated with redundant information processing. By contrast, synergy constraints were lower overall and more spatially concentrated.

### 2.3 Structure-function constraint strength (SFCS) exhibits spatial heterogeneity across the cortex

To depict the cortical distribution of constraint strength, we summed the group-level SFCS spectrum across structural components for each region, defined this measure as regional SFCS degree, and projected it onto the cortical surface (Fig. 3C). Consistent with the component-level results, redundancy exhibited a higher overall SFCS degree than synergy, together with a broader value range (13.1-18.5 vs. 10.9-12.2), reflecting greater regional heterogeneity. In contrast, synergy showed a lower overall SFCS degree and a more spatially concentrated distribution.

Within the redundancy constraint map, high-constraint regions were concentrated in bilateral visual cortex, with additional involvement of bilateral superior frontal cortex. The six highest-ranking regions were the left and right pericalcarine cortex, left cuneus, left lingual gyrus, and left and right superior frontal gyrus, indicating that redundancy-related constraints were centered on primary and secondary visual areas with additional frontal participation. The synergy constraint map showed a different spatial distribution. The six regions with the highest degree values were the left and right medial orbitofrontal cortex, left lateral orbitofrontal cortex, the left and right superior frontal gyrus and left superior parietal lobule. Relative to redundancy, synergy peaks were shifted away from visual cortex toward prefrontal and parietal association regions, with prominent involvement of orbitofrontal and superior frontal cortex and extension into superior parietal cortex. Notably, superior frontal regions appeared among the high-ranking regions for both redundancy and synergy, suggesting that this region may contribute to both interaction classes.

### 2.4 Individual SFCS predicts informant- over patient-reported cognition

To quantify the behavioral relevance of the higher-order structural constraints and test whether they encode individual differences in cognitive profiles, we developed a novel graph neural network framework, termed the structural-scaffold SFCS-GNN (sSFCS-GNN; see Methods 5.4). Crucially, this framework leverages subject-specific SFCS profiles to guide the message-passing across cortical nodes, thereby capturing nonlinear interactions between multi-scale structural scaffolding and complex behavioral phenotypes.

Fig. 4A summarizes the predictive performance (Pearson *r*) of redundancy and synergy SFCS across a broad spectrum of cognitive scales. The red and blue points represent redundancy-based and synergy-based SFCS features, respectively, and the length of the connecting lines reflects the magnitude of the predictive advantage for the dominant regime. Rather than showing a uniform advantage of one component over the other, the two exhibited partially complementary predictive ability. The strongest association for redundancy SFCS was observed for EcogSPDivatt (*r* = 0.2568*, p <* 0.001), whereas the best predictions for synergy SFCS were found for EcogSPTotal (*r* = 0.2470*, p <* 0.001) and EcogSPLang (*r* = 0.2270*, p <* 0.001). This results revealed that SFCS serves as a robust determinant of inter-individual variation across most cognitive measures.

**Fig. 4:**
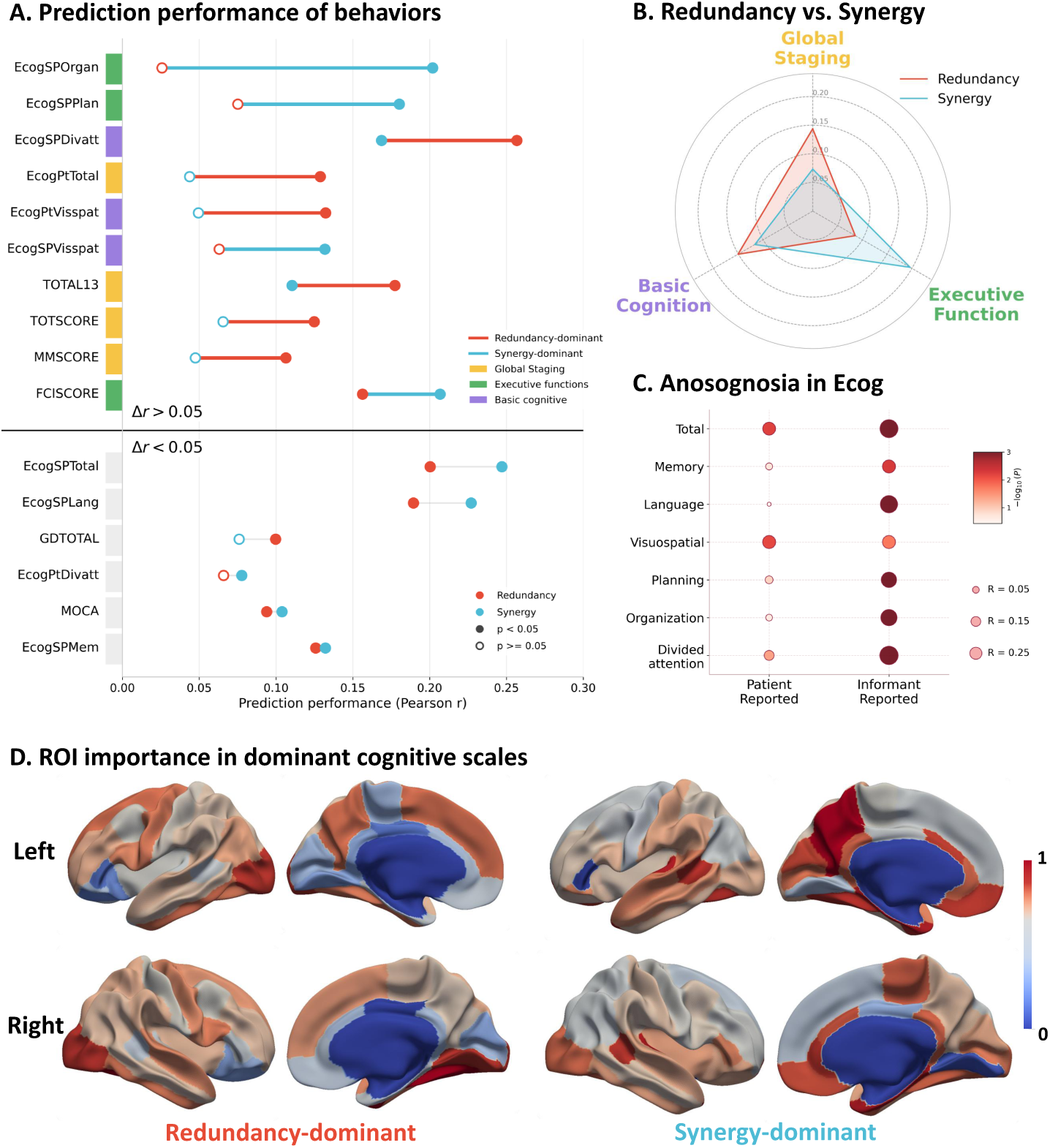
Prediction of cognitive phenotypes from individual-level redundancy and synergy SFCS features. (A) Predictive performance (Pearson *r*) across cognitive scales for redundancy (red) and synergy (blue) SFCS; (B) Radar plot showing domain-specific predictive strengths across Global Staging, Executive Function, and Basic Cognition; (C) Comparison between informant-reported and patient-reported scores across ECog subscales. Bubble size indicates *R*, and color intensity reflects − log_10_(*P* ); (D) Cortical surface mapping of sSFCS-GNN node importance for redundancy and synergy SFCS within their respective optimal predictive models.

This complementarity was further characterized across specific cognitive domains (Fig. 4B). All cognitive phenotypes were categorized into three primary domains, as indicated by the color-coded labels in the radar plot. Synergy SFCS showed a relative advantage for executive function, whereas redundancy SFCS performed better for global staging and basic cognition. Using representative examples, synergy SFCS showed stronger prediction for FCISCORE, whereas redundancy SFCS performed better for TOTAL13 and EcogSPDivatt. These results suggest that redundancy and synergy SFCS capture overlapping but non-identical information across cognitive domains.

Within the ECog measures, this predictive dissociation between redundancy and synergy was further reflected in an asymmetry between informant-reported and patient-reported assessments (Fig. 4C). Overall, SFCS predicted informant-based measures more strongly than patient-reported measures. When the larger correlation coefficient from redundancy or synergy SFCS was taken as the representative value for each subscale, all seven informant-reported subscales reached significance, with correlation coefficients ranging from 0.1318 to 0.2568 (*p <* 0.05). In contrast, only three of the seven patient-reported subscales were significant (EcogPtVisspat, EcogPtTotal, and EcogPtDivatt, *R* = 0.0777–0.1322*, p <* 0.05). This pattern suggests that SFCS-based predictions are more closely aligned with informant-reported than with patient-reported functional measures, potentially reflecting sensitivity to anosognosia, the clinical lack of insight into one’s own cognitive decline [21].

The spatial distribution of sSFCS-GNN node importance provided further anatomical context for neurobiological dissociation between redundancy and synergy constraints (Fig. 4D). The highest-contributing nodes for redundancy SFCS were concentrated in occipito-temporal visual association cortex, including the right fusiform cortex, right lateral occipital cortex, right lingual cortex, and left lateral occipital cortex; these regions are broadly linked to visual representation and perceptual processing. In contrast, the key nodes for synergy SFCS were located primarily in medial temporal and posteromedial cortices, including the left precuneus, bilateral entorhinal cortex, and right parahippocampal cortex; these regions are commonly implicated in memory, scene representation, and integration of information across regions. This clear spatial segregation confirms that redundancy and synergy SFCS capture distinct neurobiological features, each tailored to different cognitive requirements.

### 2.5 Alzheimer’s disease disrupts localized redundancy-related constraints

Having established that the SFCS profile effectively predicts cognitive abilities in healthy populations, we next sought to determine whether this structure-function constraint is disrupted in neurodegenerative conditions characterized by cognitive decline. To this end, we applied our framework to an independent cohort of AD patients (Fig. 5A). Detailed information for both groups is provided in Supplementary Table. S1.

**Fig. 5:**
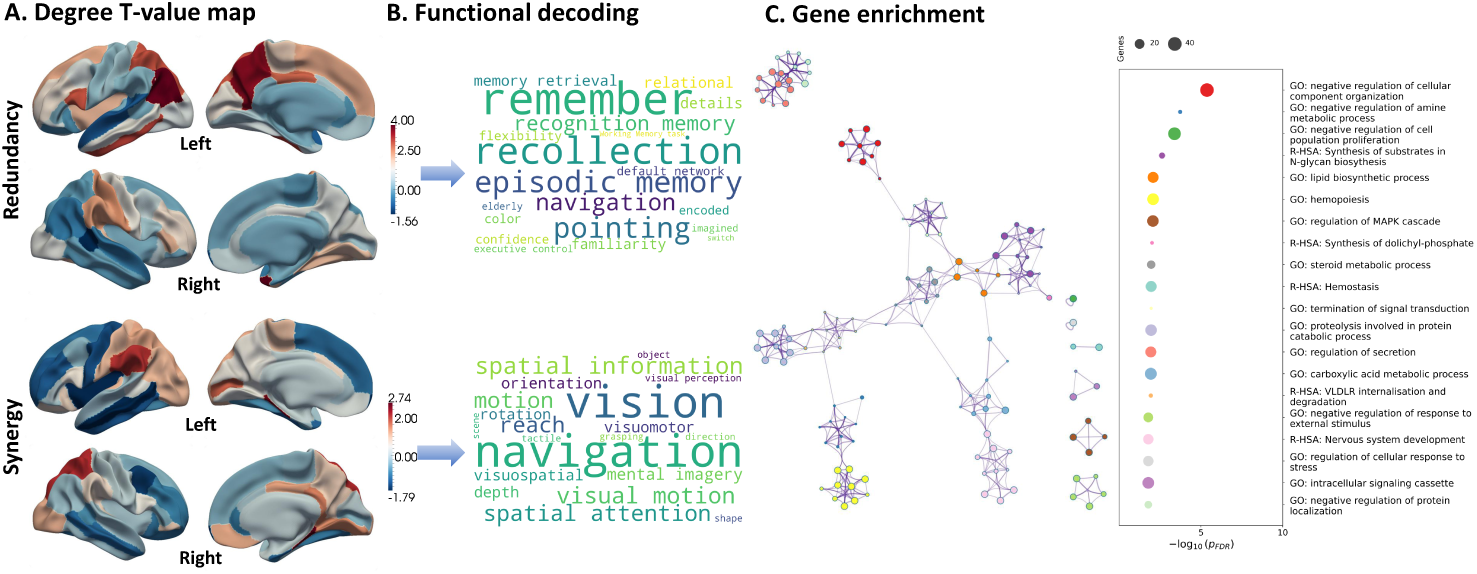
Regional SFCS degree-based network alterations in AD, together with functional decoding and gene analysis. (A) Cortical maps of group differences (AD vs. CN) in redundancy and synergy regional SFCS degree. (B) Functional decoding of constraint patterns. Word clouds visualize the neurocognitive relevance of the unthresholded T-value maps. (C) Molecular characterization of the degree T-value map, illustrating clustered biological pathways (left) and significant terms (right).

For redundancy-based regional SFCS degree, the AD group showed significantly higher values than Cognitively Normal (CN) in six regions after FDR correction: the left inferior parietal cortex, left precuneus, left superior parietal cortex, left inferior temporal gyrus, right temporal pole and right parahippocampal gyrus. These effects were concentrated in parietal and temporal association cortices, indicating that AD-related increases in redundancy-based structural constraint strength were anatomically selective rather than diffuse. By contrast, no cortical region survived FDR correction for synergy-based regional SFCS degree.

To further interpret these spatial patterns, we performed functional decoding of the unthresholded T-value maps for both redundancy-based and synergy-based regional SFCS degree (Fig. 5B). The redundancy-related map was most strongly associated with episodic-memory functions, with top terms including “remember” (*r* = 0.187), “recollection” (*r* = 0.185) and “episodic memory” (*r* = 0.162). By contrast, the synergy-related map was preferentially linked to visuospatial processing, with strongest associations for “navigation” (*r* = 0.260), “vision” (*r* = 0.257) and “spatial information” (*r* = 0.212). Thus, although synergy-related regional differences did not survive parcel-wise FDR correction, the distributed cortical pattern nevertheless aligned with a coherent visuospatial axis, whereas redundancy-related alterations preferentially tracked episodic-memory and retrieval systems.

Macroscopic brain network alterations are ultimately rooted in microscopic molecular gradients. To uncover the biological underpinnings driving the observed SFCS spatial patterns, we integrated spatial transcriptomic data from the Allen Human Brain Atlas (AHBA) [22] and performed gene enrichment analysis. Based on the T-value map of between-group differences in SFCS degree between AD and CN, we identified genes spatially associated with this regional pattern and performed functional enrichment analysis (Fig. 5C). Focusing on the leading enriched terms, the associated genes were primarily enriched for negative regulation of cellular component organization, negative regulation of amine metabolic processes, negative regulation of cell population proliferation, and synthesis of substrates in N-glycan biosynthesis. These results indicate that the regional pattern of AD-CN differences in structure-to-function constraint is preferentially associated with molecular processes related to cellular organization, metabolic regulation, cell-population homeostasis, and glycosylation-related biosynthesis.

## 3 Discussion

The structural constraint on brain function has canonically been investigated through the lens of pairwise connectivity [23, 24]. Here, we expand this paradigm by demonstrating that structural topology fundamentally orchestrates the brain’s higher-order functional landscape. We reveal that this structural constraint strength is highly heterogeneous, manifesting as stable and distinct topological patterns for redundancy-and synergy-dominant HOIs. Crucially, the derived structure-function constraint strength (SFCS) is not merely a theoretical metric, but a biologically grounded bridge across multiple scales of brain organization. Specifically, SFCS act as robust predictors of individual cognitive domains, capture targeted pathological disruptions in AD, unveil the underlying molecular signatures of disease vulnerability. Together, these findings advance the structure-function relationship from simple dyadic links to complex, biologically meaningful higher-order informational architectures.

Although functional HOIs are mathematically defined as a triplet, our findings demonstrate that they can be stably reconstructed from pairwise structural components. This indicates that higher-order functional organization does not emerge spontaneously. It is strictly tethered to underlying anatomical scaffolds [24, 25]. Pairwise structural components-capturing path reachability, global integration, and inter-regional communication connectivity do not merely yield statistical fits. Instead, they delineate the essential topological prerequisites under which complex multi-region interactions can unfold. Triadic interactions likely stabilize precisely because the participating regions are coupled through multiplexed structural representations, rather than relying on a single anatomical metric [26, 27]. More importantly, this mapping relationship still holds at the individual level, indicating that the influence of structure on higher-order functional interactions is not just a rough trend at the group-average level, but has clear individual specificity [28, 29]. This suggests that inter-individual variations in both anatomical morphology and structural connectivity serve as a crucial source of differences in higher-order functional organization.

By quantifying the anatomical constraints on higher-order functional interactions via the SFCS, we reveal a fundamental dichotomy between redundancy and synergy that unfolds across three key dimensions: overall constraint strength, structural component dependencies, and macroscopic spatial topography. First, globally, redundant HOIs exhibited stronger structural constraints than synergistic ones. This difference can be understood from the computational roles of the two types of interactions. Redundancy subserves the repeated representation of information, robust transmission, and fault-tolerant support [30, 31], and depends more heavily on anatomical pathways and communication backbones. In contrast, synergy emphasizes new information generated by the joint participation of multiple regions, and depends more on cross-system integration, dynamic state transitions, and context-dependent functional reorganization, requiring a degree of structural “untethering” to achieve cognitive flexibility [32, 33]. Second, exploring their feature dependencies reveals that communication models (CMs) and morphometric similarity networks (MSNs) provide fundamentally distinct, complementary scaffolding for these interactions. White-matter CMs characterize direct physical routing and signal propagation [26], selectively driving the heavy structural dependence of redundancy. Conversely, MSNs capture shared developmental origins and cytoarchitectural properties [34]. This morphological homophily establishes a broad, parallel structural background that facilitates functional synchro-nization—supporting both redundancy and synergy—even across regions lacking dense white matter wiring. Finally, this mechanistic asymmetry maps elegantly onto the brain’s macroscopic spatial topography along the fundamental hierarchical axis [35]. Redundant constraints securely anchor unimodal visual and sensory-related systems to ensure high-fidelity input processing. Synergistic constraints dominate transmodal frontoparietal cortices and other higher-order integrative regions, consistent with their roles in complex cognition, cross-modal integration, and flexible control [36]. Ultimately, this integrated framework highlights how the brain leverages rigid communication backbones for processing reliability, while utilizing morphological homophily to unlock higher-order cognitive flexibility.

Beyond reflecting fundamental principles of brain organization, the structure-function constraint landscape exhibits profound behavioral relevance. Crucially, redundant and synergistic SFCS do not merely encode general cognitive capacity; rather, they demonstrate a complementary predictive pattern across distinct cognitive domains, underscoring the translational potential of this quantitative framework. Specifically, synergistic constraints show a marked advantage in predicting ECogSPOrgan (representing organization, planning, and execution) [37], which reflects the strong reliance of executive functions on complex, higher-order information integration [38, 39]. Conversely, redundant constraints better predict ECogSPDivatt (involving the allocation and maintenance of attention [37]), as sustaining attentional states necessitates stable, interference-resistant, and globally coherent regulatory signals across brain networks [40, 41]. By unveiling this precise mapping among structural constraints, informational modes, and cognitive phenotypes, SFCS emerges as a highly valuable biomarker for decoding complex brain-behavior relationships. Furthermore, our predictive models exhibited greater sensitivity of informant-reported than patient-reported. This asymmetry aligns seamlessly with the clinical reality of cognitive decline: patient self-evaluations are frequently confounded by anosognosia (loss of insight) and subjective compensatory mechanisms, whereas informant reports more reliably capture sustained, externally observable functional deteriorations [42, 43]. Anatomically, the node importance weights derived from our predictive models reveal that this behavioral dissociation is rooted in a profound cortical division of labor between external perception and internal construction. Anatomically, the spatial distribution of predictive node importance reveals a profound cortical segregation between these two interaction modes. The structural constraints driving redundancy are predominantly anchored in occipito-temporal visual association cortices, aligning seamlessly with the functional demand for robust visual representation and perceptual processing. In sharp contrast, synergy-driven predictions are governed by medial temporal and posteromedial cortices—regions classically implicated in memory, scene representation, and the integration of information across distributed networks. Ultimately, this clear topographic dissociation confirms that redundancy and synergy are grounded in distinct neurobiological systems, each structurally tailored to specific cognitive requirements. By unveiling this mapping, our findings highlight SFCS as a powerful biomarker for decoding how anatomy distinctly tunes individual cognition.

In the context of neurodegeneration, the structural-functional constraint landscape undergoes striking, mode-specific reorganization in AD. Specifically, redundant constraints are pathologically intensified within memory-related hubs, whereas synergistic constraints exhibit a distributed vulnerability across visuospatial systems. Deconstructing these alterations reveals that the increase in redundant SFCS is highly anatomically selective, concentrating within the temporoparietal association cortex and medial temporal lobe—hubs canonically vulnerable to AD pathology [44, 45]. Given that redundancy subserves stable, fault-tolerant information routing [30, 31], this localized increase indicates that higher-order association regions, which typically demand flexible communication, are forced to adopt more conservative and lower-complexity strategies. This phenomenon of pathological ’redundantization’ likely reflects a restriction of flexible network dynamics, driving a rigid tethering to stable structural pathways. Remarkably, functional decoding reveals that this redundancy-related pattern preferentially tracks episodic-memory and retrieval systems, directly mirroring the signature clinical deficits of AD. In contrast, synergy-related alterations are preferentially linked to a coherent visuospatial processing and navigation axis. Although these synergistic shifts did not survive parcel-wise FDR correction, their distributed cortical pattern underscores a widespread, systemic vulnerability of complex visuospatial integration rather than a lack of disease effect. Considering the limited AD sample size, synergistic abnormalities with smaller effect sizes and more diffuse spatial distributions may be difficult to pass strict correction at the single-region level. Thus, AD is characterized by a selective reorganization of information interaction modes tailored to different cognitive axes.

Given that altered glycosylation directly impacts the degradation of AD-related proteins [46], these transcriptomic associations, paired with genetic evidence of pathways like APP degradation [47], suggest a clear molecular basis for the altered constraint topography. Importantly, this macroscopic reorganization of structural constraints is ultimately rooted in a non-random microscale molecular architecture. Spatial transcriptomic mapping demonstrates that the regional pattern of AD-CN constraint differences is preferentially coupled with genes governing negative regulation of cellular organization, amine metabolic regulation, proliferative control, and N-glycan biosynthesis. The enrichment of cellular-organization and proliferation-regulatory terms points to regional remodeling and homeostatic control, consistent with the cellular-phase framework of AD [48] and with proteomic evidence implicating metabolic, glial, MAPK/metabolism, glycosylation/endoplasmic-reticulum, and protein-transport modules in AD [49, 50]. Furthermore, the prominent link to N-glycan biosynthesis implicates disruptions in glycoprotein homeostasis, a critical pathway for protein folding, trafficking, and quality control [51]. Given that altered glycosylation directly impacts the degradation of AD-related proteins [46], these transcriptomic associations, paired with genetic evidence of pathways like APP degradation [47], suggest a clear molecular basis for the altered constraint topography. By linking these scales, our findings illustrate how microscale molecular and metabolic disruptions cascade upward to fundamentally reshape macroscopic information interaction patterns in the diseased brain.

## 4 Materials

### 4.1 Participants

The MRI scans and corresponding clinical data utilized in this study were obtained from the Alzheimer’s Disease Neuroimaging Initiative (ADNI) database [52]. As a publicly available repository, ADNI operates with approval from the Institutional Review Boards (IRB) of all participating sites. The study cohort consisted of 339 individuals (571 scans), categorized into Cognitively Normal (CN) and AD groups. All neuroimaging data included in the analysis underwent rigorous quality control procedures.

### 4.2 Preprocessing

T1-weighted images were processed with FreeSurfer v6.0 [53], including skull stripping, intensity correction and normalization, registration, and cortical reconstruction [54]. Reconstructions were visually inspected by trained researchers.Ten morphological features were extracted: number of vertices, surface area, gray matter volume, cortical thickness, thickness standard deviation [55], mean curvature, Gaussian curvature, folding index, curvature index, and sulcal depth. Feature values were mapped onto white matter surface vertices and averaged within each brain region, yielding a 10-dimensional feature vector per region.

Diffusion MRI data were preprocessed using MRtrix3 [56] with FSL [57] and FreeSurfer [53], including MP-PCA denoising, Gibbs ringing removal, and rigid-body registration to T1 space. A five-tissue-type image was generated for Anatomically Constrained Tractography (ACT). Fiber Orientation Distributions were estimated using Multi-Shell Multi-Tissue Constrained Spherical Deconvolution, followed by whole-brain probabilistic tractography using iFOD2 [58]. Streamlines were seeded from the white matter mask (FA *>* 0.2) with 100,000 selected streamlines, a maximum curvature angle of 45°, FOD cutoff of 0.05, and length range of 28–200 mm. ACT with backtracking and cropping at the gray matter-white matter interface was applied. Structural connectivity matrices were defined by the number of streamlines between brain regions.

Functional MRI data were preprocessed using SPM12 [59], FSL [57], and custom MATLAB scripts. The first 10 volumes were discarded. In SPM12, preprocessing included slice timing correction (Sinc interpolation; TR = 3.0 s), head motion correction using six-parameter rigid-body transformation, and 6-mm FWHM Gaussian smoothing. In FSL, T1-weighted images were resampled to 2-mm isotropic resolution, and the mean functional image was registered to structural space using FLIRT (6 degrees of freedom, normalized mutual information). The transformation was applied to the functional time series, followed by temporal band-pass filtering (0.01–0.1 Hz). Data were then projected onto the fsaverage6 cortical surface (41k vertices per hemisphere) using nearest-neighbor interpolation, yielding 81,924 vertices per subject [60]. Regional time series were calculated by averaging vertices within each brain region.

## 5 Methods

### 5.1 Construction of HOIs via *O*-information

To quantify higher-order statistical dependencies within macroscopic brain networks, we employed ***O***-information, an information-theoretic metric designed to delineate the dynamic balance between information redundancy and synergy in multivariate systems (e.g., sets of brain regions) [9, 61]. Mathematically, ***O***-information captures the difference between Total Correlation (TC) and Dual Total Correlation (DTC). For a set of random variables *X* = {*X*_1_*, . . . , X_n_*} representing BOLD time series extracted from *n* brain regions, the ***O***-information is defined as:

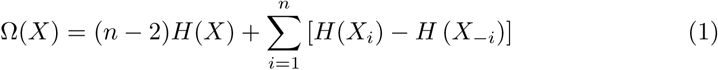

where *H*(·) is information entropy, *X*_−_*_i_* denotes the variable set excluding *X_i_*.

The sign of the ***O***-information endows the higher-order interactions with explicit physiological interpretations. A positive value (Ω *>* 0) indicates a redundancy-dominated system. In macroscopic brain networks, redundancy implies that multiple distinct regions tend to encode or transmit highly overlapping information. Redundancy is widely recognized as the foundation of the brain functional network’s robustness, fault tolerance, and resilience against focal perturbations, which are crucial for maintaining basic sensorimotor functions and the stability of local modules [62]. Conversely, a negative value (Ω *<* 0) signifies a synergy-dominated system. Synergy profoundly reflects the emergent properties of complex brain networks, wherein independent regions, acting as a dynamic and unified collective, generate novel information that strictly exceeds the linear sum of their individual contributions. Within the context of higher cognition, synergy-dominated interactions represent higher-order information integration and complex distributed computation at the whole-brain scale, serving as the core computational scaffold supporting advanced cognitive processes (such as executive control, working memory, and fluid intelligence) and adaptive cognitive flexibility [63].

Furthermore, as a model-free statistic, the computational complexity of ***O***-information scales linearly with system size, rendering it highly tractable for large-scale, whole-brain fMRI analyses. In the current study, we strictly limited the computation of ***O***-information to spatial multiplets of size *n* = 3 (triplets). This triplet-level restriction serves not only as a fundamental building block for probing higher-order topological effects but also as a crucial regularization strategy to circumvent the curse of dimensionality, which introduces substantial bias in high-dimensional entropy estimation given the limited length of fMRI time series [64, 65]. To ensure the fidelity of these higher-order network estimates, the regional blood-oxygen-level-dependent (BOLD) signals were preprocessed with Global Signal Regression (GSR) to stringently mitigate spurious covariances driven by widespread physiological artifacts and head motion. For each subject, HOIs was computed for all triplets as ***O***-information using a Gaussian copula estimator. Time series were rank-normalized to Gaussian marginals, and entropies were obtained analytically from the covariance matrix. This rigorous computational pipeline ultimately yielded a three-dimensional higher-order functional interaction matrix of shape (68 × 68 × 68) for each individual, serving as the real target variable for our subsequent structure-function constraint modeling.

### 5.2 Structural components

#### 5.2.1 Morphological Similarity Networks

To construct the morphological similarity networks (MSNs) across the 68 brain regions defined by the Desikan-Killiany (DK) atlas [66], we leveraged the 10 regional cortical morphological features extracted during the preprocessing stage. To multidimensionally and accurately quantify the covariance and similarity between any pair of brain regions across these morphological feature profiles, we systematically selected 14 distinct statistical similarity and distance metrics. The selection of these 14 methods was adapted from a comprehensive evaluation framework previously established for benchmarking mapping methods of functional connectivity in the brain (which originally evaluated 239 pairwise interaction statistics) [67]. We rigorously filtered this exhaustive repertoire to identify metrics mathematically and theoretically appropriate for comparing static, cross-regional cortical feature vectors. Based on their mathematical foundations, the final 14 selected methods were systematically categorized into four distinct classes: Rank-based Correlation Measures, Distance-based Dissimilarity Measures, Weighted Dissimilarity Measures, and Generalized Correlation Measures. The resulting 14 two-dimensional morphological similarity matrices quantitatively profile the topological associations of structural characteristics between different brain regions, serving alongside white matter communication matrices as the core multidimensional structural priors for the subsequent higher-order Structure-Function Constraint analysis.

For each subject, the morphological features were organized into a matrix *X* ∈ R*^N^*^×*F*^ , where *N* represents the number of brain regions (nodes) and *F* denotes the number of morphological features extracted per region. Let **x***_i_* = [*x_i_*_1_*, x_i_*_2_*, . . . , x_iF_* ]*^T^* and **x***_j_* = [*x_j_*_1_*, x_j_*_2_*, . . . , x_jF_* ]*^T^* be the feature vectors for region *i* and region *j*, respectively. Prior to metric calculation, feature vectors were standardized (Z-scored) across regions to ensure comparable scales. We quantified the inter-regional similarity or dissimilarity between every pair of regions (*i, j*) using the following measures.

#### Rank-based Correlation Measures

This section aims to capture the consistency of morphological trends between brain regions while mitigating the impact of outliers inherent in biological data. Since the distributions of the 10 cortical features may not strictly adhere to normality, and significant physiological variations often exist across regions, traditional linear correlations (e.g., Pearson) can be biased. Rank-based methods focus on the relative ordering of feature values rather than their absolute magnitudes, thereby providing a more robust measure of monotonic covariance in the morphological developmental profiles of brain regions.

#### Spearman’s Rank Correlation (*r_s_*)

Spearman’s *ρ*assesses how well the relationship between two variables can be described using a monotonic function [68]. It is defined as the Pearson correlation coefficient between the rank variables.

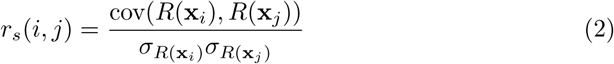

where *R*(·) denotes the rank conversion of the feature vector, cov(·) is the covariance, and *σ* represents the standard deviation of the ranks.

#### Kendall’s Rank Correlation (*τ* )

Kendall’s *τ* measures the ordinal association between two measured quantities [69]. It is defined based on the number of concordant and discordant pairs of observations.

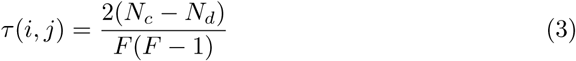

where *N_c_* is the number of concordant pairs and *N_d_* is the number of discordant pairs across the feature dimensions.

#### Connectivity Strength (Squared/Absolute)

To quantify the strength of the connection regardless of direction (positive or negative correlation), we computed the squared magnitude (or absolute value, depending on configuration) of the correlation coefficients:

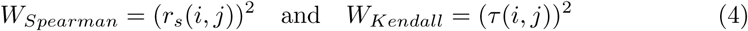

#### Distance-based Dissimilarity Measures

The purpose of this section is to quantify the proximity of brain regions within the multidimensional feature space from a geometric perspective. Treating each brain region as a point in the 10-dimensional feature space, we computed distance-based dissimilarity measures to characterize the geometric separation between regional morphological profiles.

The core measures in this category are based on the Minkowski distance, a general family of metrics that summarizes feature-wise differences between two regional vectors through an order parameter *p*. By varying *p*, this framework captures different notions of distance in the same feature space.

Minkowski Family Distances: We utilized the Minkowski distance metric across different orders *p*:

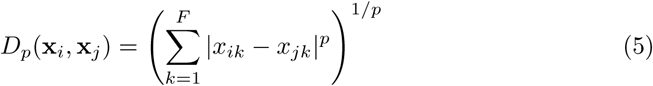

Specifically, we calculated:

**Euclidean Distance** (*L*_2_ norm): Corresponds to *p* = 2, representing the straight-line distance in the feature space [70].

**Cityblock (Manhattan) Distance** (*L*_1_ norm): Corresponds to *p* = 1, representing the sum of absolute differences [71].

**Chebyshev Distance** (*L*_∞_ norm): Defined as the limit as *p* → ∞, capturing the maximum difference along any single feature dimension [72]:

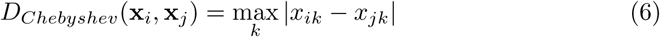

#### Cosine Distance

To measure the angular difference between feature vectors [73], ignoring magnitude, we employed Cosine distance:

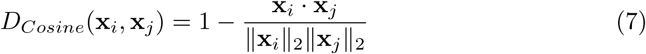

#### Weighted Dissimilarity Measures

This section aims to enhance the sensitivity of similarity measures to scale differences, particularly for features with smaller magnitudes that hold biological significance. In certain cortical features, subtle variations may reflect critical pathological or developmental information. Canberra and Bray-Curtis distances, through weighting or normalization terms, balance the contributions of different feature dimensions. This prevents features with large absolute values (e.g., surface area) from overshadowing those with smaller ranges (e.g., curvature or fractal dimension), thereby providing a more nuanced assessment of morphological dissimilarity.

#### Canberra Distance

A weighted version of the Manhattan distance, sensitive to small changes near zero [71]:

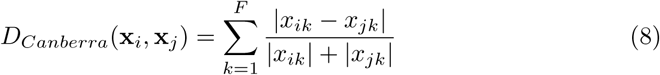

#### Bray-Curtis Dissimilarity

A normalization method often used in ecology, robust to differences in total magnitude [74]:

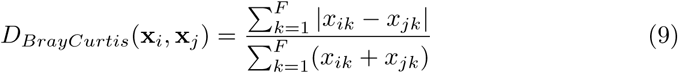

#### Generalized Correlation Measures

The objective of this section is to uncover complex, non-linear morphological coupling between brain regions. Traditional construction of morphological networks often assumes linear or monotonic relationships, which may oversimplify the biological reality. Brain development and anatomical structuring are governed by multiple non-linear factors. By employing Distance Correlation (*dCor*) and the Hilbert-Schmidt Independence Criterion (*HSIC*), we can capture arbitrary types of generalized statistical dependencies between the feature distributions of two regions without relying on specific distributional assumptions, thus revealing latent anatomical connectivity patterns that conventional methods might miss.

#### Distance Correlation (*dCor*)

Distance correlation measures both linear and non-linear association between two random vectors [75]. Unlike Pearson correlation, R(**x***_i_,* **x***_j_*) = 0 implies independence.

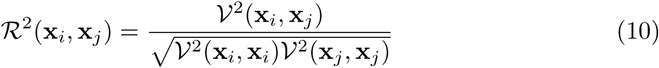

where V^2^ is the distance covariance. We computed both the standard biased estimator and the unbiased estimator (Ω*_n_*) as implemented in the dcor Python package.

#### Hilbert-Schmidt Independence Criterion (*HSIC*)

HSIC is a kernel-based independence measure [76]. We mapped the feature vectors into Reproducing Kernel Hilbert Spaces (RKHS) using a Radial Basis Function (RBF) kernel:

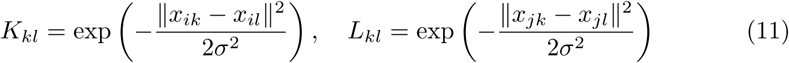

where *σ* is the kernel width (set to the median Euclidean distance). The HSIC value is computed as the normalized trace of the product of centered kernel matrices **K***_c_* and 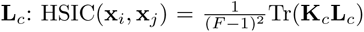 We reported both the biased estimator and an unbiased-like estimator suited for finite sample sizes.

#### 5.2.2 Communication Models

To comprehensively characterize the network communication mechanisms of white matter tracts across brain regions, we derived 9 distinct network communication models based on the structural connectivity (SC) matrices reconstructed from diffusion tensor imaging (DTI). The construction of these communication matrices was adapted from a previously established structural network analysis framework [77]. By multidimensionally decomposing the empirical SC matrices, we quantified various potential pathways and dynamic properties of signal propagation within the brain’s structural topology. Based on their underlying network routing and signaling mechanisms, these 9 communication models were systematically categorized into three broad classes: Topological Distance Measures, Topological Similarity, and Communicability and Dynamical Measures. These two-dimensional white matter communication matrices reveal the multiscale information transfer patterns of the brain network, ranging from local shortest paths to global random walks. Together with the aforementioned morphological similarity networks, they constitute a comprehensive set of multidimensional structural priors, laying the foundation for the subsequent computation of the higher-order Structure-Function Constraint Strength (SFCS).

Let *W* be the weighted adjacency matrix where *W_ij_* represents the connection strength (streamline count) between region *i* and region *j*, and *N* be the number of regions.

#### Topological Distance Measures

This section aims to quantify the global communication efficiency and integration capacity within the brain network. In DTI-derived structural networks, connection weights reflect the integrity or density of white matter tracts; higher weights are hypothesized to facilitate faster signal propagation. Consequently, by mapping inverse weights to topological distances, we identify the most economical routes for information transfer between regions. These metrics are fundamental for characterizing neurobiological signal latencies and identifying backbone pathways that enable the rapid integration of distributed information.

#### Weighted Shortest Path Length (*SPL*) and *Hops*

To characterize the efficiency of information transmission, we computed the shortest paths based on the inverse of the connection weights. The mapping from weight to length was defined as *L_ij_* = 1*/W_ij_*, under the assumption that higher connection strength facilitates shorter communication latency. The weighted shortest path length 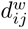 between node *i* and node *j* was calculated using the Floyd-Warshall algorithm:

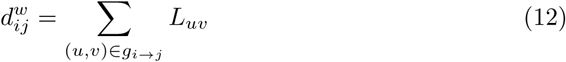

where *g_i_*_→*j*_ is the geodesic path minimizing the total weighted distance. SPL refers to the matrix containing 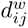 for all pairs. Simultaneously, we recorded the number of hops (*h_ij_*), representing the number of edges constituting the shortest path. The algorithm also generated a predecessor matrix (Pmat), which stores the sequence of nodes along the shortest path, used for path reconstruction.

#### Binary Distance (*BD*)

Complementary to the weighted metrics, we computed the binary distance on the binarized adjacency matrix *A* (where *A_ij_*= 1 if *W_ij_ >* 0, and 0 otherwise). *D_bin_*(*i, j*) represents the minimum number of edges required to traverse from node *i* to node *j* without considering edge weights.

#### Topological Similarity

Relying solely on the shortest path assumption may not fully capture complex neural signaling patterns, as biological networks often utilize redundant connections to ensure robustness. The purpose of this section is to evaluate the local contextual architecture and network transitivity. By computing topological similarity and path transitivity, we assess whether shortest paths are supported by dense local neighborhoods (i.e., the availability of local detours). High transitivity typically indicates a structural backbone that is resilient to perturbation and conducive to functional segregation and specialized processing.

#### Matching Index (*MI*)

To quantify the similarity of conn ectivity profiles between two regions, we calculated the Matching Index [78]. For any pair of nodes *i* and *j*, *M_ij_* measures the overlap of their neighbors, excluding any direct connection between them:

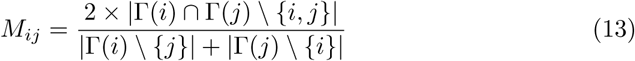

where Γ(*i*) denotes the set of neighbors of node *i*. This index ranges from 0 (no common neighbors) to 1 (identical connectivity profiles).

#### Path Transitivity (*PT* )

We employed Path Transitivity to assess the density of local detours available along the shortest paths. It is defined as the average Matching Index of all consecutive node pairs involved in the shortest path between *i* and *j*:

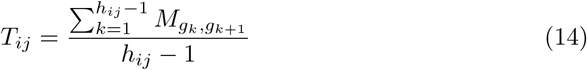

where *g_k_* is the *k*-th node in the shortest path sequence *g_i_*_→*j*_. High path transitivity indicates that the shortest path is supported by a dense local neighborhood structure [79].

#### Communicability and Dynamical Measures

Neural communication is not restricted to single optimal trajectories; signals often propagate across the whole-brain network via parallel channels in a diffusive manner. To capture these dynamic interactions, we employed metrics based on random walks and diffusion processes. Unlike distance-based measures, communicability and mean first passage time account for all possible walks between nodes—including non-shortest paths—thereby providing a more comprehensive characterization of how structural connectivity supports large-scale neural dynamics and functional coupling.

#### Communicability (*F* )

To capture the ease of communication between nodes considering all possible walks (not just the shortest ones), we computed the weighted communicability [80]. The matrix was first normalized to account for node strength differences: 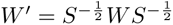, where *S* is the diagonal matrix of node strengths. The communicability matrix *F* is defined using the matrix exponential:

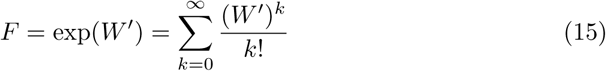

where the (*i, j*)-th entry accounts for the total contribution of walks of all lengths between *i* and *j*, weighted by their factorial decay.

#### Mean First Passage Time (*MFPT* )

We modeled a discrete-time random walk on the network to compute the Mean First Passage Time [81]. The transition probability matrix is defined as

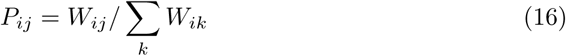

The MFPT, denoted as *H_ij_*, represents the expected number of steps a random walker takes to reach node *j* for the first time starting from node *i*. It is derived from the fundamental matrix *Z* = (*I* − *P* + Π)^−1^, where Π is a matrix with rows equal to the stationary distribution vector 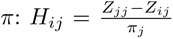 This metric quantifies the “diffusive distance” between regions.

#### Flow Graph Dynamics (*FGD*)

To assess the global information flow over continuous time, we simulated a continuous-time random walk (CTRW) process. The dynamics are governed by the graph Laplacian L. The flow graph at a specific Markov time *t* (set to *t* = 1 in our analysis) is given by the propagator of the density:

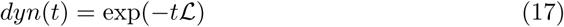

The resulting matrix captures the probability density of random walkers redistributing over the network after time *t*, reflecting the temporal scale of integration [82].

### 5.3 Quantification of Structural-Functional Constraint Strength (SFCS)

To quantify the extent to which structural architecture constrained higher-order functional interactions, we implemented a structure–function regression framework. All analyses were performed on the 68 cortical regions defined by the Desikan-Killiany (DK) atlas [66]. For each participant, a triadic HOI tensor (68×68×68) was computed from resting-state fMRI signals, with each element representing the ***O***-information of a given triplet of brain regions. According to the sign of ***O***-information, higher-order interactions were classified as redundancy-dominated (Ω *>* 0) or synergy-dominated (Ω *<* 0). To focus on the strongest higher-order effects, we extracted the upper 20% and lower 20% of the HOIs distribution for each participant as target signals and modeled them separately.

Structural predictors consisted of 23 two-dimensional matrices, including 9 communication models and 14 morphological similarity networks. For each participant and each HOI target set, slice-wise fitting was performed separately for each region. Specifically, for a given region *k*, the upper-triangular elements of the corresponding two-dimensional HOI slice were vectorized and used as the dependent variable, and the upper-triangular elements from the matched locations of the 23 structural matrices were vectorized and used as predictors in a generalized linear model [83] with an intercept term. For region *k*, the model is

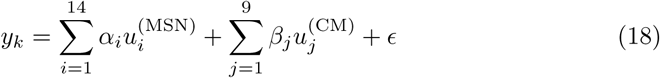

where *y_k_* is the vectorized upper-triangular elements of the HOI slice corresponding to brain region *k*, 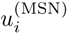 denotes the vectorized upper-triangular elements of the *i*-th two-dimensional MSN matrix, 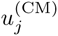 denotes the vectorized upper-triangular elements of the *j*-th two-dimensional CM matrix, *α_i_* and *β_j_* denote the corresponding weight coefficients for MSN and CM predictors, respectively, and *ɛ* is the error term.

This procedure yielded, for each participant and for each HOI target set, a coefficient matrix of shape 68 × 24, in which the first column corresponded to the intercept and the remaining 23 columns corresponded to the structural-component coefficients. In the present study, structure–function constraint strength (SFCS) was defined as the 68 × 23 coefficient matrix obtained after removing the intercept column. The SFCS derived from fitting the upper 20% of the HOIs distribution was defined as redundancy SFCS, whereas the SFCS derived from fitting the lower 20% of the HOIs distribution was defined as synergy SFCS. These SFCS features were used in downstream group comparisons and biological interpretation analyses.

### 5.4 Cognitive Prediction via sSFCS-GNN

We developed the structural-scaffold SFCS-GNN (sSFCS-GNN) to predict individual cognitive recognition scores from SFCS features embedded in sample-specific structural brain graphs. As illustrated in Fig. 6, each sample was represented as an input brain graph, in which the 68 ROIs were treated as graph nodes, sample-specific structural connections were used as graph edges, and SFCS values were assigned as node features. Specifically, each ROI was represented by a 23-dimensional SFCS feature vector, resulting in 68 nodes with 23 features per node for each sample. The sample-specific structural connectivity matrix provided the graph adjacency among the 68 ROIs.

**Fig. 6:**
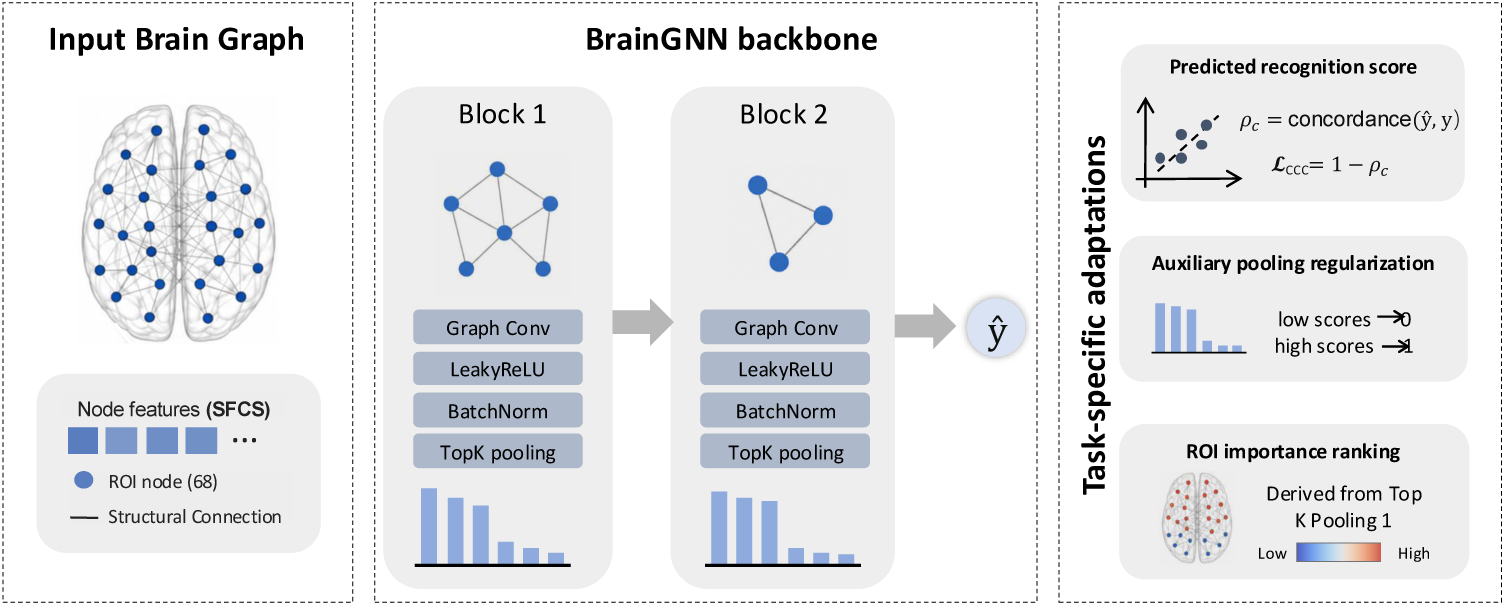
Overview of the sSFCS-GNN framework. Individual structural brain graphs, with 68 ROI nodes and SFCS-derived node features, are processed by a two-block BrainGNN backbone to predict recognition scores. Task-specific adaptations include CCC-based optimization, pooling regularization, and ROI importance ranking from the first TopK pooling layer.

The model comprised two hierarchical graph-learning blocks followed by a regression head. Each block consisted of a graph convolution layer, LeakyReLU activation, batch normalization, and TopK pooling with a pooling ratio of 0.5. After the second pooling stage, node representations were flattened and passed through a multilayer perceptron to predict a continuous cognitive score *y*^. In addition, the node scores from the first pooling layer were retained for ROI importance estimation.

Model optimization was primarily driven by the concordance correlation coefficient (CCC) loss:

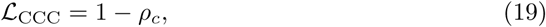

where

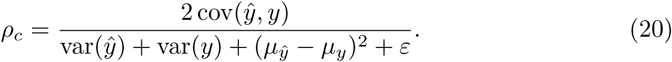

Here, *µ_y_*_^_ and *µ_y_* denote the means of the predicted and observed scores, respectively, and *ε* is a small constant introduced for numerical stability. This loss was chosen because it captures not only linear association but also agreement in location and scale between predicted and observed values.

To stabilize hierarchical node selection during pooling, the training objective further included regularization terms associated with the pooling operation. The total loss was defined as

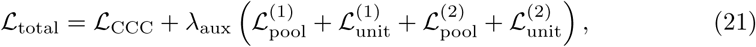

with *λ*_aux_ = 0.1. For each pooling layer, let **s** ∈ [0, 1]*^N^* denote the sigmoid-normalized node scores and let *k* = ⌊*rN* ⌋ with pooling ratio *r* = 0.5. After sorting scores in ascending order, the pooling regularization term was defined as

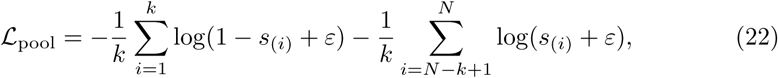

which encourages low-ranked scores to approach 0 and high-ranked scores to approach 1. The scorer weight vector **w** was additionally regularized as

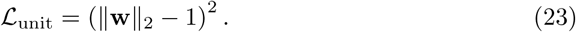

A separate regression model was trained for each cognitive measure within a cross-validation framework. Predictive performance was quantified by aggregating out-of-sample predictions across test folds and computing the Pearson correlation coefficient *r* and its parametric *P* value against the observed cognitive scores. Statistical significance was further assessed using permutation testing, in which cognitive labels were randomly shuffled 1,000 times and the same modeling procedure was repeated to generate a null distribution of correlation coefficients. A permutation-based *P* value was then computed to assess whether the observed predictive performance exceeded chance level.

To improve interpretability, ROI importance was estimated from the node scores produced by the first pooling layer. For each fold, first-layer pooling scores were averaged across test subjects, and the final ROI importance was obtained by averaging these fold-wise values across folds:

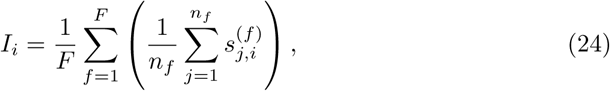

where *I_i_* denotes the final importance of ROI *i*, *F* is the number of folds, *n_f_* is the number of test subjects in fold *f* , and 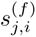 is the first-layer pooling score of ROI *i* for subject *j* in fold *f* .

### 5.5 Characterizing SFCS Alterations in AD

Group differences in regional constrained degree between patients with AD and CN were tested using inverse probability weighting (IPW) [84, 85]. Imaging-derived regional measures were matched to demographic records, and only participants with complete age and sex data were included. Propensity scores were estimated using logistic regression with diagnostic group as the dependent variable and age and sex as predictors. Stabilized IPW weights were then derived. To reduce the impact of extreme values, propensity scores were bounded to 0.001–0.999 and weights were truncated at the 1st and 99th percentiles. Covariate balance before and after weighting was assessed using standardized mean differences, and effective sample size was calculated.

Group comparisons were performed at the ROI level. For each of the 68 regions, weighted least squares regression was fitted with regional constrained degree as the dependent variable and diagnostic group as the predictor, using IPW-derived weights. The regression coefficient for group, together with its *t* statistic and two-sided *P* value, was used to quantify the AD-versus-CN effect. Multiple comparisons were controlled using the Benjamini–Hochberg false discovery rate procedure, with FDR-corrected *P <* 0.05 considered significant.

To functionally characterize the regional-level statistical map, we performed meta-analytic decoding using the BrainStat toolbox [86]. Following the standard BrainStat pipeline, the input map was spatially correlated with term-specific meta-analytic activation maps precomputed within the Neurosynth framework, and cognitive terms were ranked according to their Pearson correlation coefficients with the input map [87]. The complete decoding results were exported as a table for further inspection. In addition, positively correlated terms were visualized using a word cloud to provide an intuitive summary of the dominant cognitive associations represented by the statistical map.

### 5.6 Gene expression decoding of degree T-value

To determine whether the spatial distribution of degree T-value was related to regional transcriptional variation, we performed imaging-transcriptomic decoding using partial least squares (PLS) regression. Redundancy-related regional SFCS degree values across the 68 cortical parcels of the Desikan–Killiany atlas were used as the imaging phenotype, and matched parcel-wise cortical gene expression profiles were obtained from the Allen Human Brain Atlas using abagen [88]. Genes with missing expression values across all parcels were excluded, remaining missing values were imputed using the across-parcel mean for each gene, and both the gene expression matrix and the regional SFCS degree vector were z-scored across parcels. After preprocessing, 15,633 genes were retained for analysis.

The primary PLS analysis focused on the first latent component (PLS1), which captures the dominant mode of covariation between cortical gene expression and the spatial pattern of SFCS degree. This primary component captured a substantial proportion of spatial variance (*R*^2^ = 0.384; Fig. S4A), substantially dominating the remaining latent dimensions (Fig. S4B). To assess weight stability, we performed 5,000 bootstrap resamples across cortical parcels with replacement, with bootstrap solutions sign-aligned to the original PLS1 solution at each iteration. Bootstrap *Z*-scores were calculated from the mean and standard deviation of the resampled weights, converted to two-sided *P* -values, and corrected for multiple comparisons using the Benjamini–Hochberg false discovery rate procedure.

Because the aim of this analysis was to characterize transcriptional signatures associated with higher SFCS degree, downstream enrichment analyses were restricted a priori to genes with positive PLS1 weights that survived false discovery rate correction (FDR *<* 0.05). This statistical thresholding successfully isolated 1,685 positively associated genes and 1,334 negatively associated genes along the PLS1 weight spectrum (Fig. S4C). Of the positively associated genes, the top 600 with the highest bootstrap *Z*-scores were submitted to Metascape for gene enrichment analysis [89] using the default human background gene set.

## 6 Supporting information

**Table S1:** Demographic and scan characteristics of the CN and AD groups. Values are reported at the scan level unless otherwise indicated. Age is presented as mean ± standard deviation, with median and interquartile range in parentheses.

| Characteristic | CN | AD |
| --- | --- | --- |
| Number of scans | 510 | 61 |
| Number of participants | 301 | 38 |
| Female, n (%) | 315 (61.76%) | 16 (26.23%) |
| Male, n (%) | 195 (38.24%) | 45 (73.77%) |
| Age, years | 73.80 ± 8.10 | 75.95 ± 7.22 |
| Age, median (IQR) | 73.00 (68.00–79.00) | 78.00 (71.00–81.00) |
| Age range, years | 56.00–95.00 | 55.00–88.00 |
| Scan years | 2017–2023 | 2017–2022 |

**Fig. S1:**
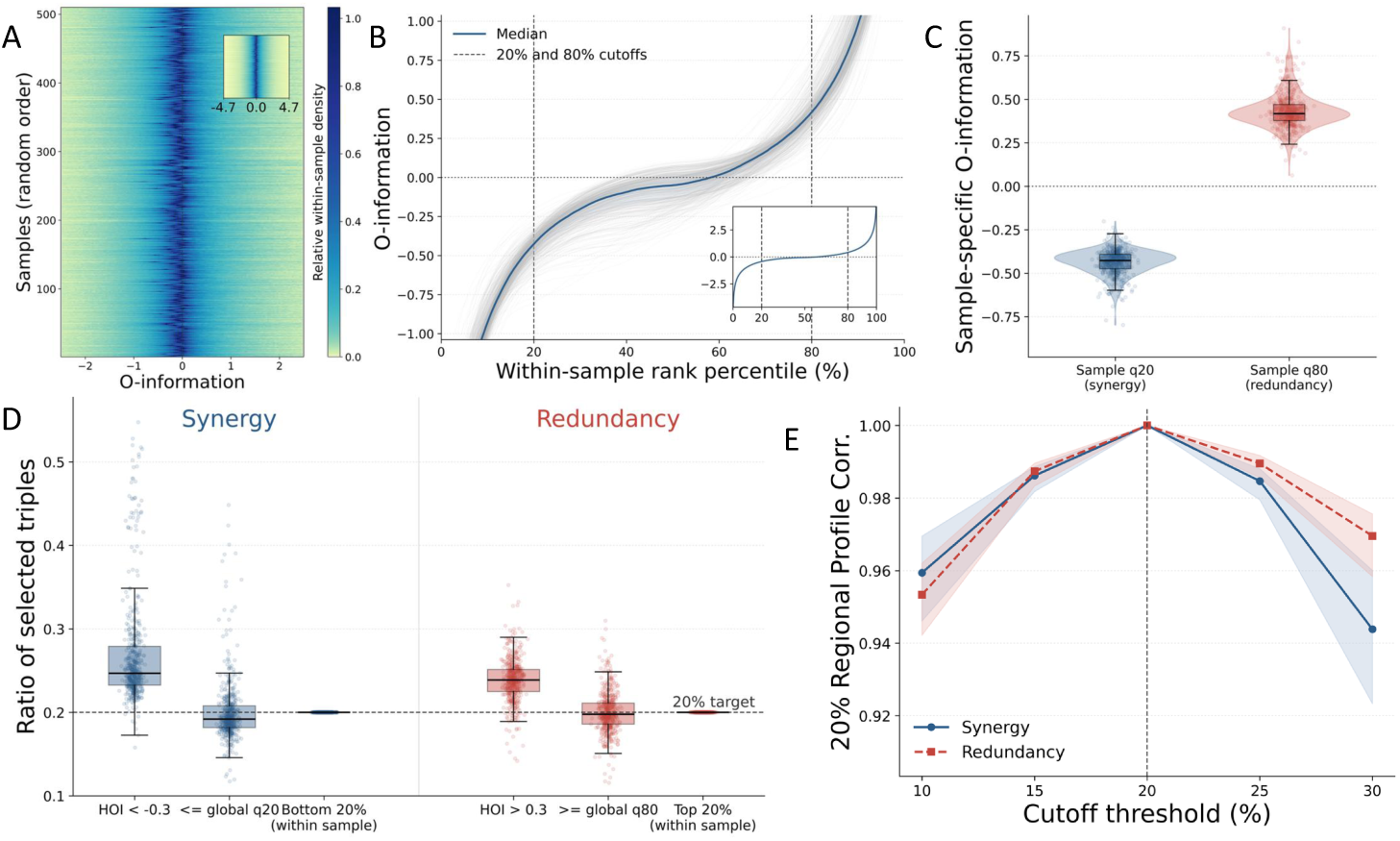
Supplementary experiment supporting the rationale for applying a 20% threshold. (A) Heatmap of *O*-information values across all unique DK68 triplets (*i < j < k*) for each subject. Subjects are shown in random order; the main panel focuses on the central range (−2 to 2), and the inset shows the full range. (B) HOIs values were ranked within subject from lowest to highest. Gray lines indicate individual subjects, the blue line indicates the median, and shading indicates the interquartile range; dashed lines mark the 20% and 80% cutoffs. Most values form a central plateau near 0, whereas large-magnitude negative and positive values are concentrated in the tails. (C) Distribution of the within-subject *q*_20_ and *q*_80_ thresholds across all subjects. *q*_20_ lies in the negative range and *q*_80_ in the positive range, but both vary substantially between subjects, indicating that equivalent tail definitions do not correspond to a single absolute HOIs value. (D) Fraction of selected triplets per subject under different thresholding strategies. Absolute thresholds (HOI *<* −0.3, HOI *>* 0.3, or global *q*_20_*/q*_80_) produce marked between-subject variability, whereas within-subject *Bottom 20%* and *Top 20%* retain a stable selection fraction of 20% for every subject. (E) Correlation between regional participation profiles obtained at different tail cutoffs (10%–30%) and the profile obtained at 20%. Profiles remain highly similar around 20%, supporting the robustness of this cutoff choice.

**Fig. S2:**
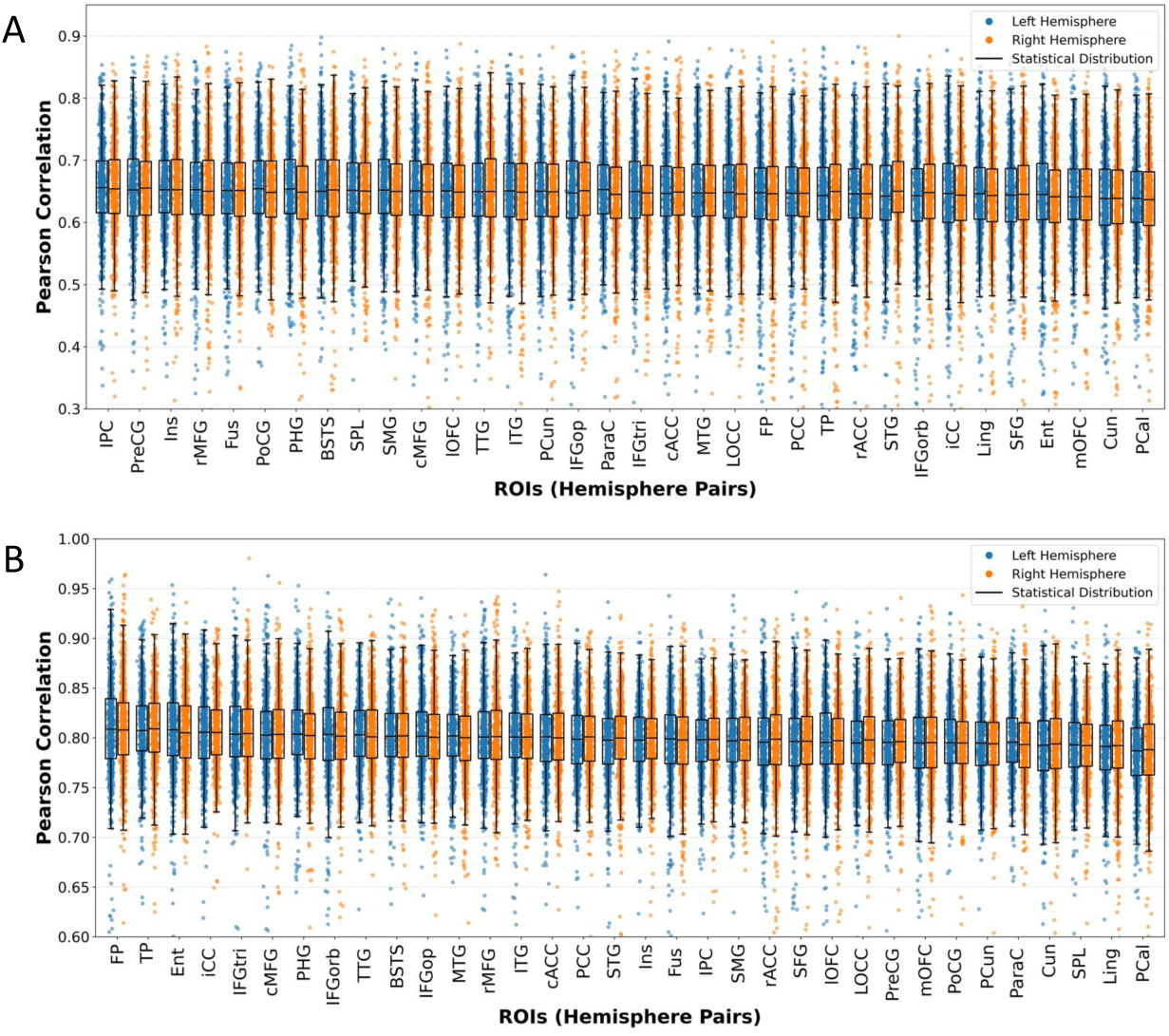
Reconstruction performance of structural components on functional HOIs under different thresholding criteria. (A) shows the reconstruction performance without thresholding. (B) shows the reconstruction performance after applying a threshold of ±0.3.

**Fig. S3:**
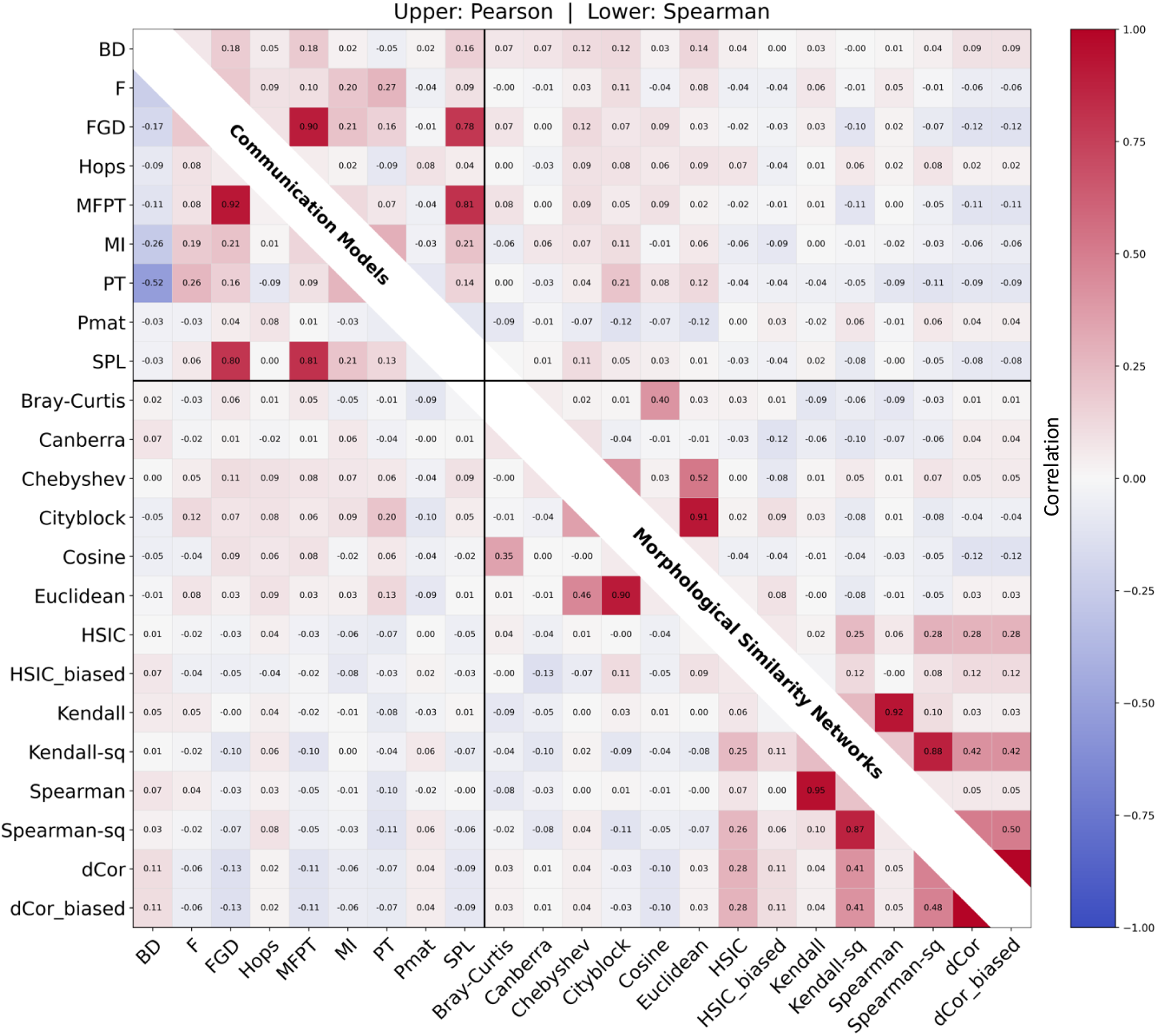
Correlation among the 23 structural components at the group level. The upper triangle shows Pearson correlations, and the lower triangle shows Spearman correlations.

**Fig. S4:**
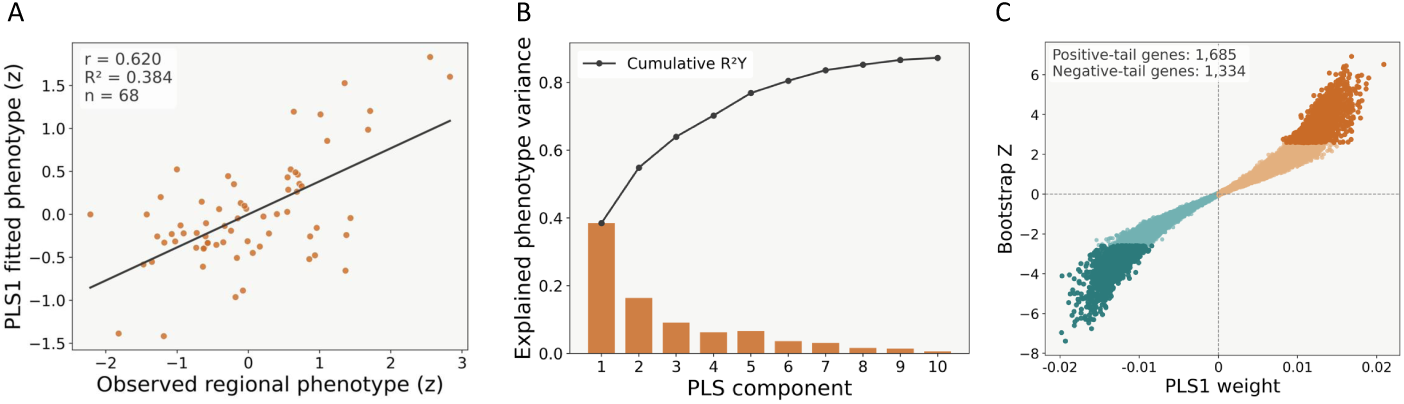
PLS decoding of the AD–CN SFCS degree T-value map. PLS regression was used to associate the regional SFCS degree T-value map across 68 cortical parcels with Allen Human Brain Atlas gene expression profiles. (A) Observed versus PLS1-fitted phenotype (*r* = 0.620, *R*^2^ = 0.384). (B) Variance explained by the first ten PLS components. (C) PLS1 gene weights and bootstrap *z* scores, highlighting significant positive- and negative-tail genes.

## Acknowledgements

Songyao Zhang was supported by the Young Scientists Fund of the National Natural Science Foundation of China (No. 62403103), Joint Funds of the National Natural Science Fundation of China (No. U24A20753), Dalian Municipal Guiding Program for the Life and Health Sector (No. 2025ZDJH01PT040) and the Fundamental Research Funds for the Central Universities (No. DUT25YG272). Marco Palombo was supported by the UKRI Future Leaders Fellowship MR/T020296/2 and 1073, as well as the MRC Research Grant MR/031566/1. Xi Jiang was supported by the National Natural Science Foundation of China (62276050, 62576077). Tuo Zhang was supported by the National Natural Science Foundation of China (62476222, 62131009). Hongkai Wang was supported by the Joint Funds of the National Natural Science Fundation of China (No. U24A20753), the general program of National Natural Science Fund of China (No. 81971693), the funding of Liaoning Key Lab of IC & BME System and Dalian Engineering Research Center for Artificial Intelligence in Medical Imaging and Liaoning technology innovation center of hyperpolarized MRI.

## Data availability

The dataset analyzed during the current study was obtained from the Alzheimer’s Disease Neuroimaging Initiative (ADNI) database (https://adni.loni.usc.edu/). ADNI imaging and clinical data are available to qualified researchers through the ADNI data access process after registration and approval of the ADNI data use agreement. No ADNI individual-level imaging or clinical data are redistributed with this study.

## Code availability

The custom code used to perform the main analyses and generate the figures in this study is available at https://github.com/shichaosu/SFCS. Higher-order functional interactions (HOIs) and ***O***-information computations were performed using the publicly available hoi package (https://brainets.github.io/hoi/).

